# Ex-situ cultivation of Isoetes cangae and Isoetes serracarajensis (Isoetaceae) two endemic species from Brazilian Amazon

**DOI:** 10.1101/861351

**Authors:** Daniel Basílio Zandonadi, Rodrigo Lemes Martins, Luis Alfredo dos Santos Prado, Heitor Monteiro Duarte, Mirella Pupo Santos, Emiliano Calderon, Ana Carolina Almeida Fernandes, Quézia Souza Santos, Filipe Junior Gonçalves Nunes, Luis Carlos Felisberto Ribeiro, Taís Nogueira Fernandes, Alexandre Castilho, Francisco de Assis Esteves

## Abstract

*Isoetes* L. is a genus of lycophytes widely distributed around the world that has a large number of endemic species. Here we document the first successful large scale *ex-situ* cultivation of the new endemic species from Brazilian Amazon quillworts *Isoetes cangae* and *Isoetes serracarajensis*. These isoetids are endemic of an iron mining site and grow on a superficial iron crust that occurs over ferriferous rocks. This study aimed to develop the cultivation methods of the threatened species *I. cangae* and monitoring its unique physiology. Plants from both species brought from Amazon lagoons were cultivated through a year in a low-cost system at a different site during different seasons. The survival rate of plants was higher as 98% and both species developed well under cultivation but showed different patterns during linear growth: *I. cangae* showed faster leaf development but was slower on sprout production than *I. serracarajensis*. The mechanism of leaf expansion is related to plasma membrane H^+^-ATPase activation, near to 2-fold higher in *I. cangae*. On the other hand, the effective quantum yield of photosystem II was higher in *I. serracarajensis* than in *I. cangae*. During the cultivation, new sporophytes of *I. cangae* were produced, confirming its reproductive status. We have also tested elevated iron levels on the growth of plants, but no interference of iron concentration was observed. The results of this work have broad applicability, assisting other low-cost cultivation studies, which are very important in ecosystem recovery of mining areas and conservation strategies.

## 1. INTRODUCTION

The Isoetaceae family is composed of a single genus, *Isoetes* L., which comprises about 350 species (James Hickey et al., 2006). They are mainly aquatic or semi-aquatic of oligotrophic lagoons and slow streams, although some species grow in wet soil that dries in the summer. In Brazil, 25 species of *Isoetes* were identified (Pereira et al., 2016; Prado et al., 2015). Recently, two new endemic species in Brazilian Amazon were described: *Isoetes cangae and Isoetes serracarajensis* (Pereira et al., 2016; Santos et al., 2019). These two species distinguish from each other and other Amazon Basin species in traits of the megaspores (female spores) (Pereira et al., 2016), usually considered in this group phylogeny. However, the more interesting aspect about these species is the habitat occurrence of these two species in Serra de Carajás, Pará state, more specifically in ferruginous plateaus of the Carajás mountain range. This environment presents extremely conditions such as acid and oligotrophic soil with high concentrations of heavy metals, high temperatures, and strong seasonality, with a well-defined dry season (Gagen et al., 2019). *I. cangae* is an aquatic plant endemic to only one lagoon in the ferruginous fields of the south of Serra dos Carajás while *I. serracarajensis* is an amphibious species found in several ferruginous plateaus, in seasonally flooded environments.

The ontogeny of *I. cangae* plants and a protocol for reproduction was studied based on *in vitro* culture of megaspores and microspores in order to obtain viable sporelings (Caldeira et al., 2019). However, knowledge about the species traits and strategies related to habitat occupation, growth, survival, and reproduction in this harsh environment is fundamental for the conservation biology of *Isoetes*, and site restoration strategies.

In the present study, we aimed to establish the *ex-situ* cultivation of both *I. cangae* and *I. serracarajensis* while we explore and compare their physiological aspects related to nutrition, growth, and photosynthesis. We were able to establish a successful protocol for the outdoors’s *ex-situ* growth for both species. Besides, we provide some insights and discussion about physiological traits, which can be used as markers for ecophysiological performance for these Isoetids, such as the chlorophyll fluorescence and the plasma membrane H^+^-ATPase activity.

## 2. MATERIAL AND METHODS

### 2.1. Plant Site and Material

The two *Isoetes* species studied were endemic from Serra dos Carajás, in the southeast of the Amazon region and the Pará State, northern Brazil. Serra dos Carajás comprises north and south ranges (Serra Norte and Serra Sul, respectively), located above 700 m in altitude. Two conservation units were created by the Federal Government of Brazil to protect the Carajás mountain range: the Carajás National Forest (FLONA) and the National Park of “Campos Ferruginosos” (Ferruginous Fields). These protected areas include extensive areas of canga (superficial iron crust that occurs over ferriferous rocks) that hosts several endemic plant species just recently studied in exquisite detail (Nunes et al., 2018; Santos et al. 2019). Plants of *I. cangae* and *I. serracarajensis* were collected in the summer of 2018 (February), from different sites, according to Figure 1, to represent all the genetic and morphological diversity found in the FLONA. *I. cangae* is restricted to one lagoon called Amendoim (6°23’ 51.81” S – 50°22 20.88” W) while *I. serracarajensis* has a spread distribution along several pounds at ferruginous plateaus of Carajás Mountain range (Figure 1).

**Figure 1.**
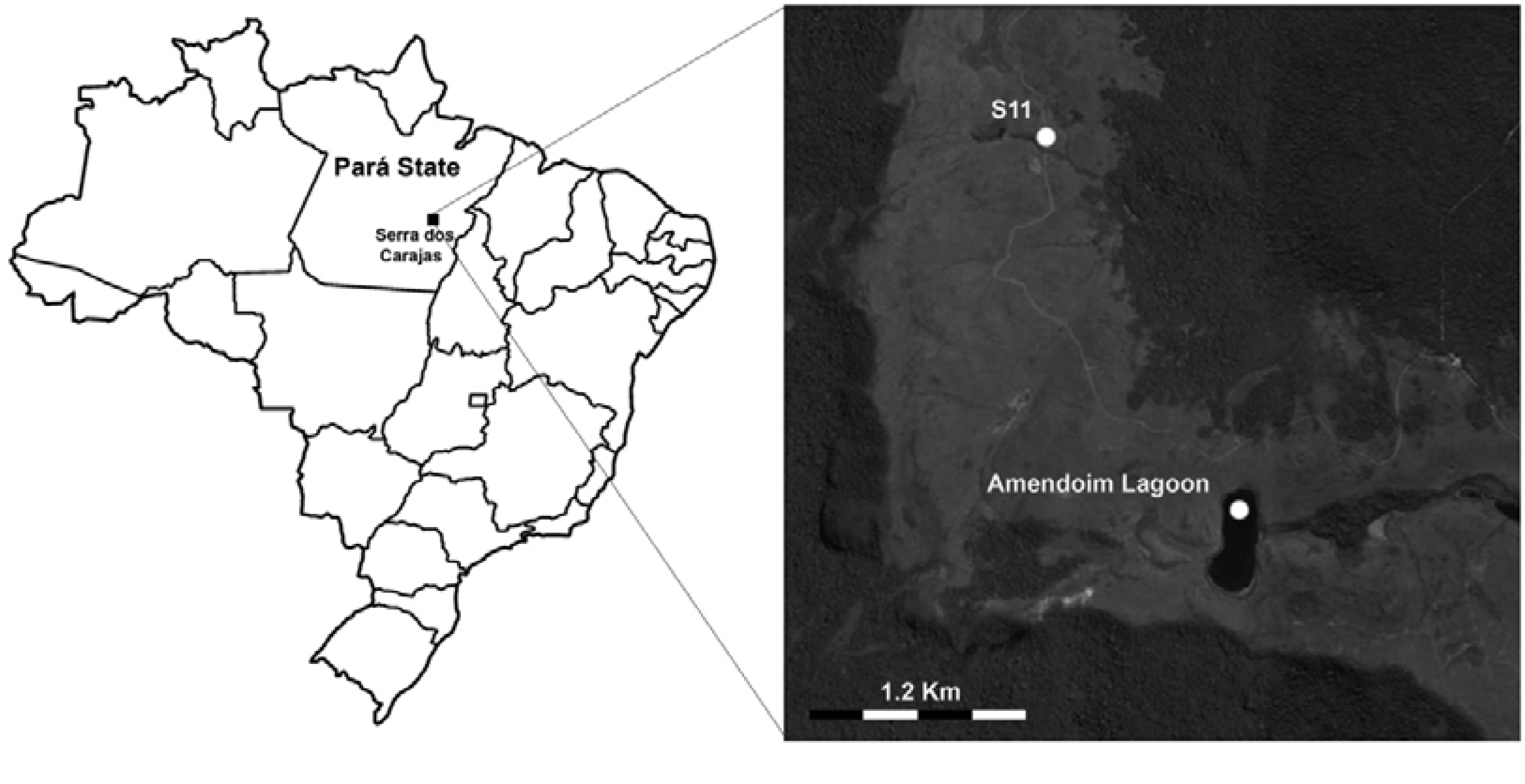
Iron Rocky Outcrops from Serra dos Carajás, Pará State, Brazil. *I. cangae* was collected in Amendoin lagoon, while *I. serracarajensis* was gathered in the ferruginous plateaus, flooded S11.

Vouchers of studied populations can be found at the Herbarium of the Federal University of Rio de Janeiro, RFA (RFA 41140 to 41142, *I. cangae*, and RFA 41143). Collecting permits were granted by Instituto Chico Mendes de Biodiversidade do Ministério do Meio Ambiente (ICMBio/MMA; n. 59724).

### 2.2. Water and Soil Analysis

Sediment samples for nutrient determination were taken in 2018 during both wet and dry seasons. During campaigns, we also take water for nutrient determination at a depth of 0.5m in the lagoon and at the pond water surface where we found *I. serracarajensis*. The following parameters were analyzed in both sediments and water samples: pH, organic matter (OM), total N (nitrogen), P (phosphorus), K (potassium) and total Fe (iron), Zn (zinc), Mn (manganese), and Cu (copper). Organic matter was determined using the method described by Walkley and Black (1934), total N was determined by the Kjeldahl method and total P by the colorimetric method (Yoshida et al., 1971). The concentrations of K, Mn, and Cu in tailings were determined by digestion with concentrated HNO_3_ and HClO_4_ (3:1) and analyzed through Inductively Coupled Plasma Emission Spectroscopy (ICPE-9000, Shimadzu).

### 2.3. Plant *ex-situ* Cultivation

Plants of both species were collected in the natural habitat and transferred to a greenhouse for monitoring the plant growth and the parameters of water quality in the new artificial environment. After collect, plants were washed to remove the substrate from the original collection sites then were conditioned in plastic pots with tap water before transporting to the laboratory in NUPEM (22°19′38.06’’;041°44′13.16’’) - Macaé City, Rio de Janeiro State, Brazil).

In laboratory *I. cangae* and *I. serracarajensis* were placed in thirty (30) previously prepared plastic beakers (2L) containing fifty milliliters of tap water, 6.21g of organic substrate (sphagnum peat) in the bottom and 21g of quartzite sand on top. The chemical parameters of the substrate are presented in table 1. Fifty (50) *I. cangae* specimens were distributed in 25 beakers, two plants in each. Ten (10) specimens of *I. serracarajensis* were placed in five (5) beakers similarly prepared the set of *I cangae*. The experimental sets of beakers were kept in the greenhouse with natural light conditions.

**Table 1.** Water characteristics of original sites of *Isoetes cangae* and *Isoetes serracarajensis*

| Species |  | *I. serracarajensis* | *I. cangae* |
| --- | --- | --- | --- |
| Location |  | Pond | Lake |
| N-NH<sub>4</sub><sup>+</sup> | mg/L | 1.00 | 1.00 |
| N-NO<sub>3</sub><sup>-</sup> | mg/L | 0.00 | 0.00 |
| B | mg/L | 0.06 | 0.06 |
| Al | mg/L | 1.84 | 2.16 |
| EC | dS/m | 0.06 | 0.01 |
| N-total | mg/L | 0.00 | 0.00 |
| P | mg/L | 0.22 | 0.20 |
| K | mg/L | 18.00 | 0.15 |
| Ca | mg/L | 0.20 | 0.20 |
| Mg | mg/L | 0.04 | 0.02 |
| Na | mg/L | 0.23 | 0.39 |
| Cu | mg/L | 0.01 | 0.01 |
| Fe | mg/L | 1.30 | 0.55 |
| Mn | mg/L | 0.10 | 0.10 |
| Zn | mg/L | 0.01 | 0.01 |
| S | mg/L | 1.67 | 1.66 |
| pH | - | 6.30 | 5.60 |

### 2.4. Plant Pruning and Growth Monitoring

Plant specimens of similar size were selected for the cultivation experiment. At the beginning of the experiment, the specimens were pruned to ensure better standardization and to facilitate the detection and measurement of new leaves.

Afterward, a regular monitoring protocol was carried out every three days during 40 days in 3 seasons (autumn, winter, and spring). After each period of 40 days, plants were pruned again, and the regrowth was monitored. Plants were evaluated daily for general conditions and growth of microorganisms. Monitoring protocol considers the registration of new leaves by counting and length measurements. Besides, we recorded plant conditions with a photographic apparatus, controlling focal distance, exposure, speed, and other photographic parameters.

The water physicochemical characteristics (pH, temperature, conductivity, salinity, dissolved solids content, percentage of oxygen saturation, and dissolved oxygen rate) and climate parameters (temperature, humidity, and luminosity) were also recorded.

### 2.5. Plant Nutrients Analysis

All leaves pruned were used to evaluate the concentrations of nutrients through inductively coupled plasma spectrometry. *Isoetes* leaves were harvested, washed, drying, and digested with concentrated HNO_3_ and HClO_4_ (3:1) to be analyzed. The analyses were performed using an Inductively Coupled Plasma Emission Spectroscopy (ICPE-9000, Shimadzu).

### 2.6. Plant Tolerance to Iron

To evaluate the *I. cangae* and *I. serracarajensis* tolerance to different concentrations of soluble iron, we exposed both to four different concentrations of Fe-EDTA added in the water (0.00 - control, 0.20, 0.60 and 1.80 mg / L). The concentrations were used based on natural total iron concentration *in-situ* (table 2). This experiment was carried out in 16 beakers, 4 per treatment, assembled similarly to the beakers used to evaluate the growing pattern. Each beaker contained three plants. Measurements of growth and sprouting and assessments of water characteristics were carried out every seven days. The results presented are representative from one season (end of winter - August).

**Table 2.**
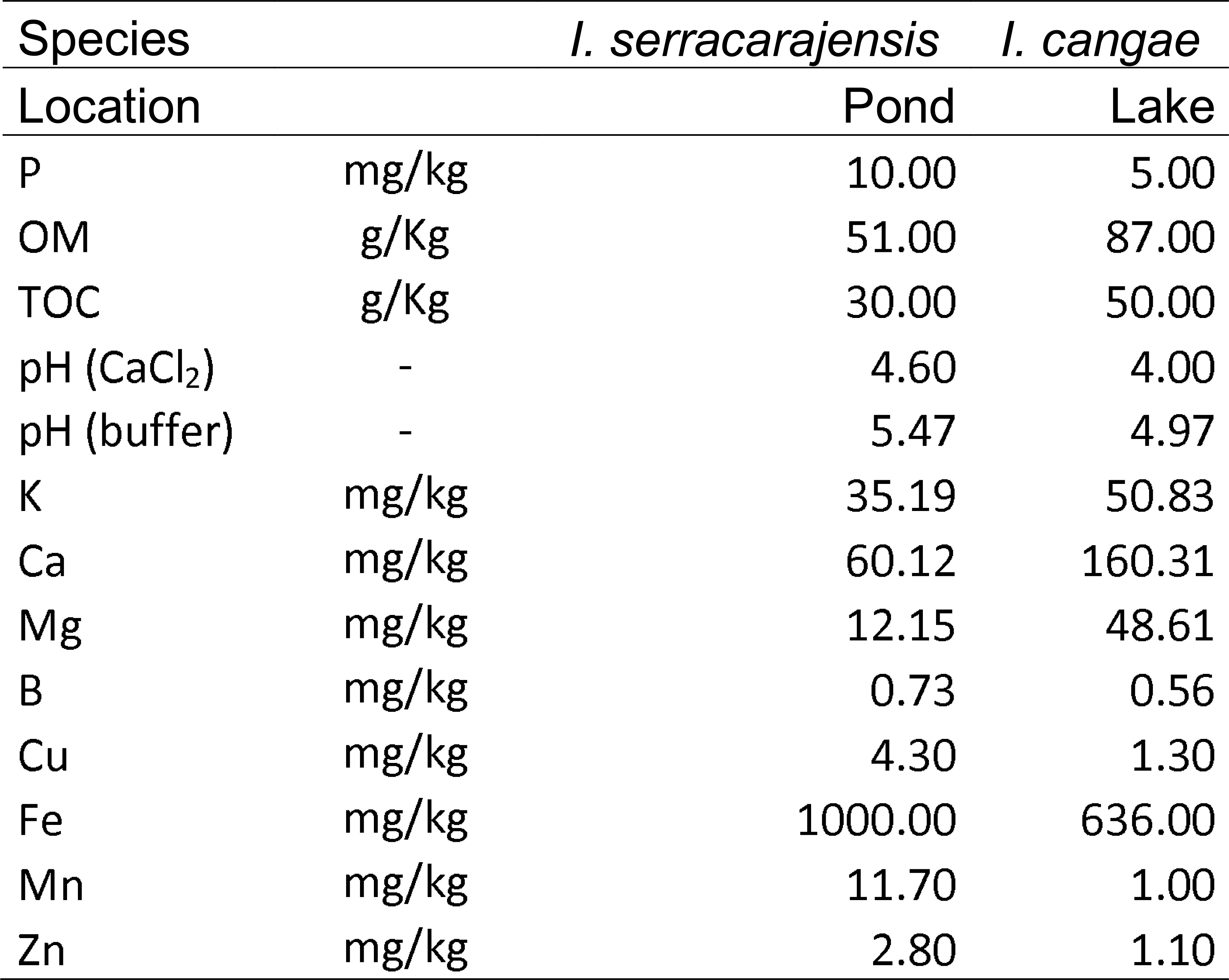
Substrate characteristics of original sites of *Isoetes cangae* (lagoon) and *Isoetes serracarajensis* (pond).

| Species |  | *I. serracarajensis* | *I. cangae* |
| --- | --- | --- | --- |
| Location |  | Pond | Lake |
| P | mg/kg | 10.00 | 5.00 |
| OM | g/Kg | 51.00 | 87.00 |
| TOC | g/Kg | 30.00 | 50.00 |
| pH (CaCl<sub>2</sub>) | - | 4.60 | 4.00 |
| pH (buffer) | - | 5.47 | 4.97 |
| K | mg/kg | 35.19 | 50.83 |
| Ca | mg/kg | 60.12 | 160.31 |
| Mg | mg/kg | 12.15 | 48.61 |
| B | mg/kg | 0.73 | 0.56 |
| Cu | mg/kg | 4.30 | 1.30 |
| Fe | mg/kg | 1000.00 | 636.00 |
| Mn | mg/kg | 11.70 | 1.00 |
| Zn | mg/kg | 2.80 | 1.10 |

### 2.7. Plasma membrane-enriched vesicles

Plasma membrane (PM) vesicles were isolated from *Isoetes* leaves using differential centrifugation as described by (Zandonadi et al., 2010) with some modifications. Briefly, about 5 g (fresh weight) of *Isoetes* leaves was homogenized using a mortar and pestle in 6 mL of ice-cold buffer containing 250 mM sucrose, 10% (w/v) glycerol, 0.5% (w/v) PVP (PVP-40, 40 kDa), 2 mM EDTA, 0.2% (w/v) BSA, and 0.1 M Tris–HCl buffer, pH 7.6. Just before use, 150 mM KCl, 2 mM DTT, and 1 mM PMSF were added to the buffer. The homogenate was strained through four layers of cheesecloth and centrifuged at 1,500*g* for 10 min. The supernatant was centrifuged at 8,000*g* for 10 min and then at 100,000*g* for 40 min. The pellet was resuspended in a small volume of ice-cold buffer containing 10 mM MES–BTP, pH 7.5, 10% (v/v) glycerol, 1 mM DTT, and 1 mM EGTA. The vesicles were either used immediately or frozen under liquid N_2_ and stored at −80°C until use. All procedures were carried out below 4°C. Protein concentrations were determined by the method of (Bradford, 1976).

### 2.8. Plasma membrane H^+^-ATPase activity

The hydrolytic H^+^-ATPase activity in PM vesicles was determined by measuring the release of Pi colorimetrically as described in Zandonadi et al. (2010). In *Isoetes cangae* plants between 70 and 80% of the PM vesicle H^+^-ATPase activity measured at pH 6.5, was inhibited by 0.1 mM sodium orthovanadate (Na_3_VO_4_), a very effective inhibitor of PM H^+^-ATPase. In *Isoetes serracarajensis* plants, only 30-40% of the PM vesicle H^+^-ATPase activity was inhibited by Na_3_VO_4_. The assay medium consisted of 1 mM A TP–BTP, 5 mM MgSO_4_, 10 mM MOPS–BTP (pH 6.5), 100 mM KCl, 0.2 mM Na_2_MoO_4_ and 0.05 mg mL^-1^ vesicle protein.

### 2.9. Chlorophyll *a* fluorescence

#### 2.9.1. Pulse-Amplitude-Modulated (PAM) fluorometry

In order to compare the photosynthetic performance of the two studied *Isoetes* species regarding leaf exposition to air, the effective quantum yield of photosystem II (Δ*F*/*Fm’,* or PSII efficiency) was accessed with a portable pulse amplitude fluorometer (mini-PAM, Walz, Germany). The photosynthetic performance of the two studied *Isoetes* species consider two points in the same leaf, below and above the waterline. The leaves were previously exposed to an irradiation 160 μmol photons m^-2^ s^-1^ provided by the mini-PAM’s internal halogen lamp, which was plugged to power line during the measurements to keep the light intensity constant. This process avoids variations on the light microenvironment. The steady-state fluorescence, recorded at ambient light (*F*) and the maximal fluorescence (*Fm’*), recorded after a saturating light pulse (800, PAR > 4000 μmol photons m^-2^ s^-1^), were used to calculate Δ*F*/*Fm’*, (where Δ*F* = *Fm*’-*F*) (Genty et al., 1989; Schreiber et al., 1995). Light intensity was recorded using the quantum sensor localized at the mini-PAM’s leaf-clip.

#### 2.9.2. Chlorophyll a fluorescence Imaging

Images of the effective quantum yield of photosystem II (Φ_PSII_) were undertaken using an imaging system developed to work as described in (Duarte et al., 2005; Duarte and Lüttge, 2007; Rascher et al., 2000). Four arrays of 36 blue light provided the photosynthetic and excitation lights-emitting diodes (λ = 470 nm, maximum power 432 Watts), the intensity of which was micro-controlled by pulse-width modulation at a frequency of 1200 Hz. During the experiments, the light intensity was kept constant at 200 μmol photons m^-2^ s^-1^. Chlorophyll fluorescence was selectively detected by a Peltier-cooled digital camera Alta U6 (Apogee Inc, USA), equipped with a CCD sensor of 1024×1024 pixels and 16-bit digitalization. A λ < 665 filter RG-655 Schott (Mainz, Germany) was attached to the camera objective (60 mm macro lens, Nikkor, USA). Images of chlorophyll fluorescence were recorded and processed on a PC by customized software written in Visual C^++^. Φ_PSII_ was recorded according to the saturating light pulse method (Genty et al., 1989; Schreiber et al., 1995). First, an image of the sample steady-state fluorescence (*iF*) under constant light intensity was recorded. After that, a saturating light pulse (intensity: ∼3000 μmol photons m^-2^ s^-1^, duration: 800 ms) was applied over the algal samples. The last 150 ms of this pulse was used to integrate the maximal fluorescence signal of the second image (*iFm*). Both *iF* and *iFm* (integer 16-bitts images) were corrected by dividing them by the pixel mean of a fluorescence standard (Walz, Germany) placed inside the image field. The resulting images (32-bits real point flow) were used to calculate the images of *i*Φ*_PSII_* were recorded every 20 min and calculated as *i*Φ*_PSII_* = (*iFm* e *iF*) / *iFm*. Posterior image processing for pixel average of individual sample and temporal dynamic was conducted with the software ImageJ (Abramoff et al., 2004).

### 2.10. Statistical analysis

Data were analyzed by ANOVA and Dunnett’ S test to determine the differences between treatments and controls. Regression analysis was performed on data of leaf growth and sprouting. For all statistical tests, p-values < 0.05 are considered statistically significant.

## 3. RESULTS

During the growth period of the *Isoetes* plants, in autumn, winter, and spring, the air temperature was, on average, 23.3 **°**C, 23.8 **°**C, and 24.8 **°**C, respectively. The relative humidity was, on average, 78.6 %, 73.4%, and 77.4%, respectively.

The water characteristics of the lagoon of *Isoetes cangae* and the pond of *Isoetes serracarajensis* are presented in table 1. Ponds and the studied lagoon distinguish in Boron (B), Aluminum (Al), Electric conductivity (EC), Phosphorous (P), Potassium (K), Magnesium (Mg), Ferrous (Fe), Sulfur (S), and Potential hydrogenionic (pH). The substratum characteristics from the original sites of each species are were distinct for all variables considered (Table 2).

Both species growth and reproduce in an artificial media developed. The cost of the cultivation system was around US$ 0.15 per plant, considering the following materials: sand, sphagnum peat, and beakers. The chemical characteristics of the organic substrate used in the cultivation of *Isoetes cangae* and *Isoetes serracarajensis* are showed in Table 3, with values of Organic Matter, Total N, Total Organic Carbon and macro and micronutrients (NH^+^_4_, P, K, Ca, Mg, S, B, Cu, Fe, Mn, Zn). Plant nutrition was accessed by quantifying leaves mineral concentration in both species (Table 4). *I*. *cangae* from Serra dos Carajás presented higher N, P, S, Cu, Fe, Mn, and Al than plants from *ex-situ* cultivation. Comparing *I. cangae* and *I. serracarajensis*, the later always presented enhanced mineral content as compared with the former.

**Table 3.**
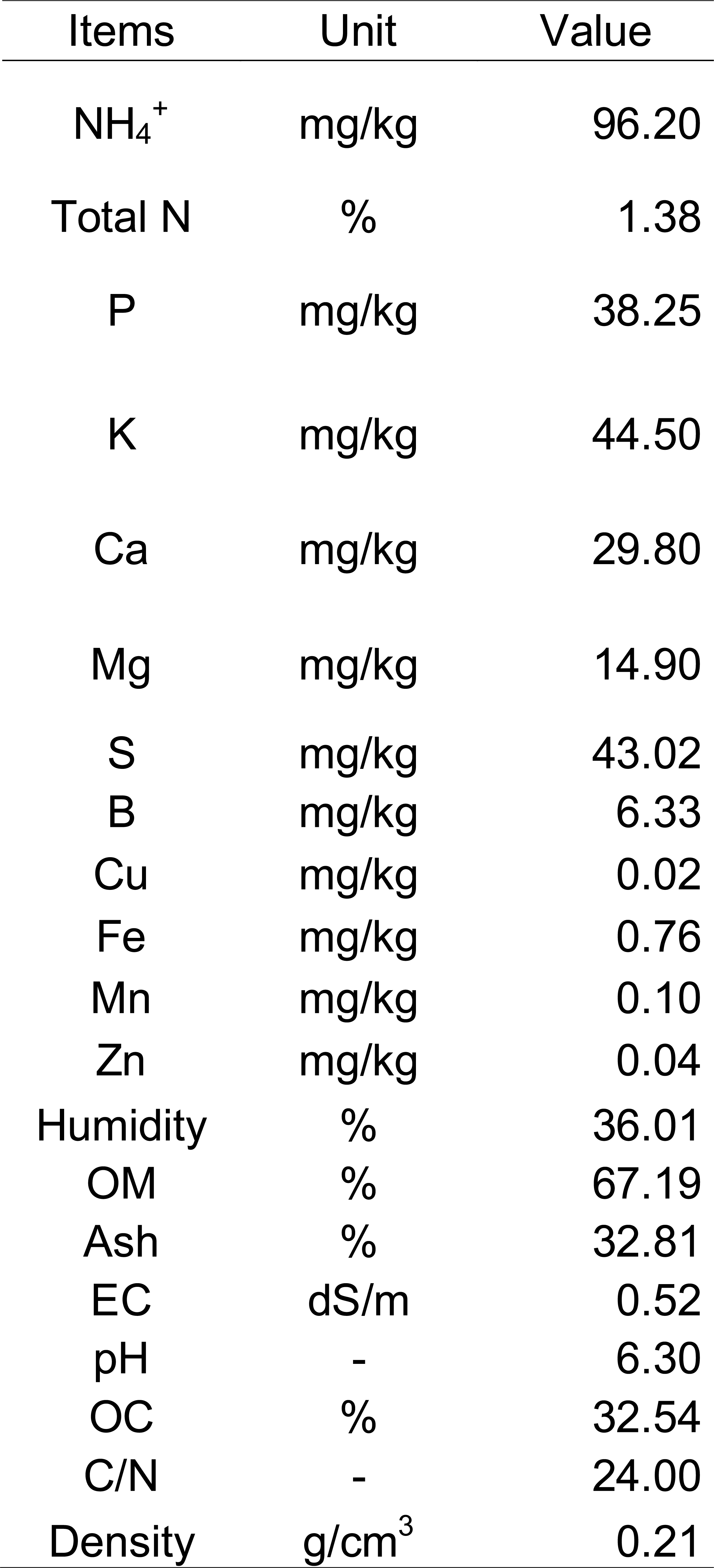
Substrate characteristics of *ex situ* cultivation medium of *Isoetes cangae* and *Isoetes serracarajensis*.

**Table 4.** Leaf tissue mineral concentrations of *Isoetes cangae* and *Isoetes serracarajensis* in two sites of Serra dos Carajás, or in *ex-situ* cultivation.

| Species |  | *I. serracarajensis* | *I. cangae* | *I. serracarajensis* | *I. cangae* |
| --- | --- | --- | --- | --- | --- |
| Location |  | Pond | Lake | *Ex-situ* cultivation |  |
| N | g/kg | 32.55 | 28.73 | nd | 19.42 |
| P | g/kg | 2.28 | 1.97 | nd | 1.34 |
| K | g/kg | 46.00 | 10.10 | nd | 13.75 |
| Ca | g/kg | 3.75 | 3.00 | nd | 9.00 |
| Mg | g/kg | 2.83 | 1.89 | nd | 2.95 |
| S | g/kg | 2.45 | 2.45 | nd | 0.84 |
| B | mg/kg | 51.87 | nd | nd | 97.76 |
| Cu | mg/kg | 15.86 | 9.60 | nd | 4.50 |
| Fe | mg/kg | 33,050.00 | 16,950.00 | nd | 150.00 |
| Mn | mg/kg | 322.95 | 135.65 | nd | 120.00 |
| Al | mg/kg | 1,080.00 | 980.00 | nd | 184.15 |
| Zn | mg/kg | 78.00 | 40.75 | nd | 76.00 |

The water physicochemical characteristics in the beakers during the analyzed period remained very similar between the two species of *Isoetes*. The temperature dropped gradually from the third day when it was 27 °C reaching the minimum point on the 12th day, with 20 °C for both species. After the 15^th^ day, temperature values increase until they return to 27°C for *I. cangae* and reaching 26 °C in *I. serracarajensis* water (Figure 2). The pH in the beckers was slightly more acidic in *I. cangae* (pH 6.58) than in *I. serracarajensis* (pH 6.92) at the beginning of the monitoring, but the values showed an abrupt increase in the sixth day (7.49 in *I. cangae* and 7.90 in *I. serracarajensis*) (Figure 3). From 12th day values were slightly higher in *I. cangae* (7.60) than in *I. serracarajensis* and later remained stable until the end of observations, on the 18th day. Conductivity showed a typical pattern in evaluated species, showing a gradual increase until the ninth day, when the conductivity was higher in *I. cangae*, falling on the 12th day when the conductivity becomes higher in *I. serracarajensis* (Figure 2). The values were stable at the end of the observations. Salinity did not change throughout the evaluation period, maintained at 0.1 ppt. Total dissolved solids had a gradual increase in both species until the 12th day; however, on the 15th day, these values decreases a little, mainly *I. cangae* (Figure 2). The levels of dissolved O_2_ and O_2_ saturation in water are quite different about the two species. *I. cangae* varied from 7.1 mg/L and 89.4% saturation on the third day, reaching 9.59 mg/L and 118.4% at the end of the experiment. *I. serracarajensis* presented values of 8.63 mg/L and 108.18% of saturation on the third day, ranging up to 9.23 mg/L and 113.58% saturation on day 8.

**Figure 2.**
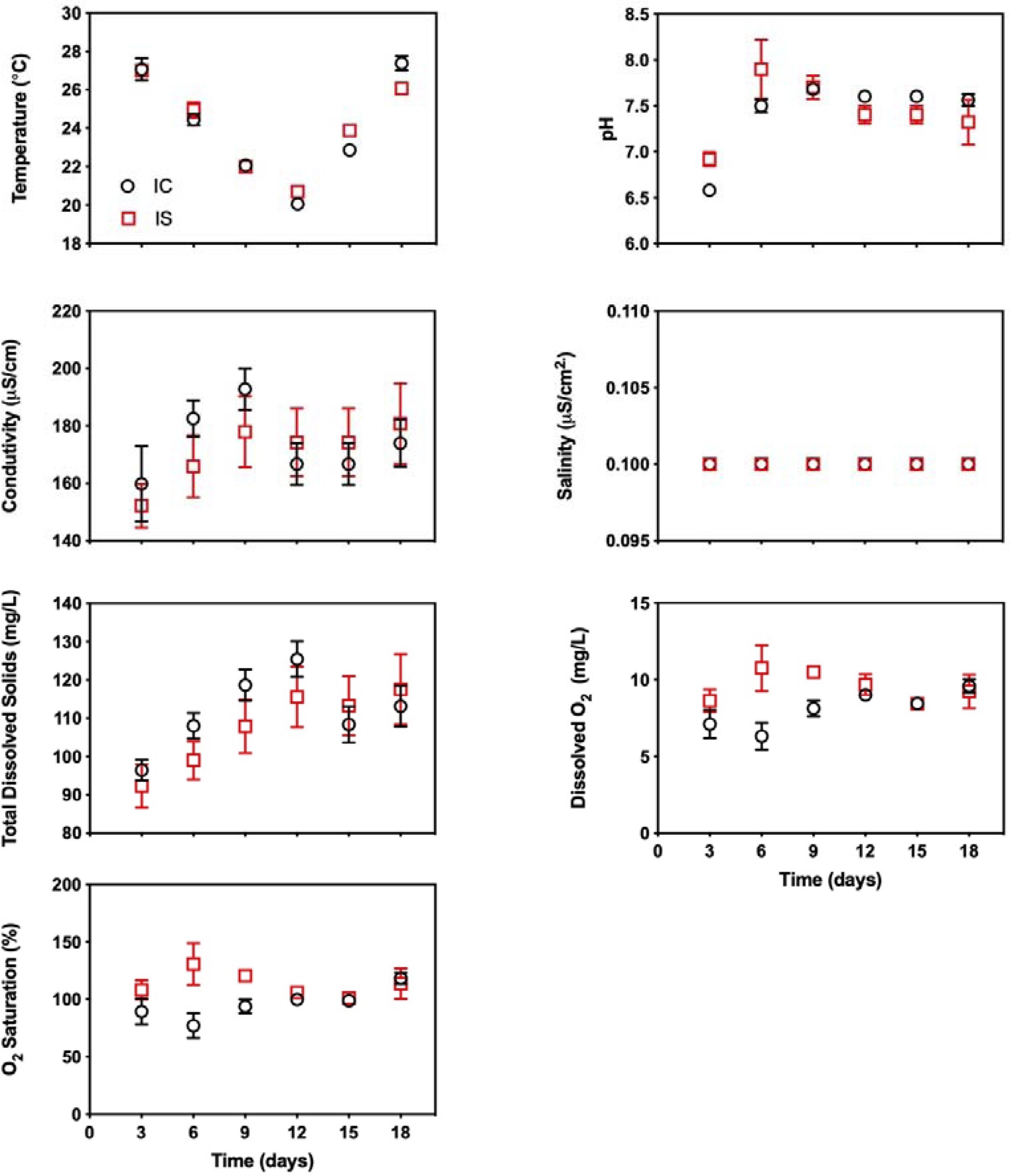
Physicochemical characteristics of the water. Temperature, pH, Conductivity, Salinity, Total Dissolved Solids, Dissolved O_2_ and O_2_ Saturation. *Isoetes cangae* (I.C. – open black circles) and *Isoetes serracarajensis* (I.S. – open red squares).

**Figure 3.**
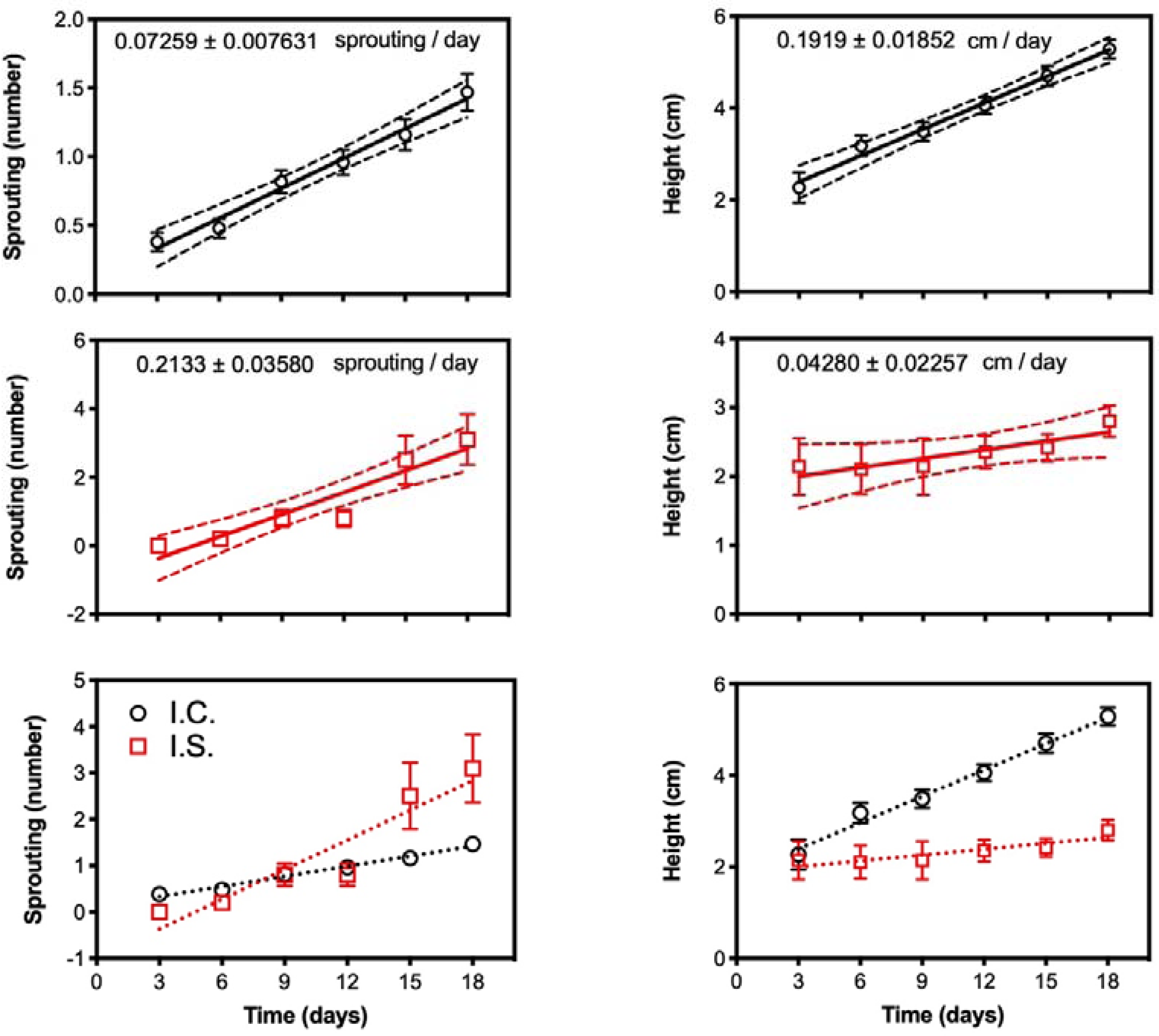
Sprouting rate in number of new leaves, growth in centimeters and comparative charts between species of *Isoetes cangae* (I.C. – open black circles) and *Isoetes serracarajensis* (I.S. – open red squares) in beakers. Data represent means from three independent experiments ± confidence interval (n= 12 in each experiment).

Plants of *I. serracarajensis* had a lower sprouting rate as compared to *I. cangae* in the first six days (Figure 2). The rates are similar on days 9, and 12 days, however *I. serracarajensis* has a budding rate approximately three times higher than *I. cangae* in the final evaluation period. An inverse pattern was observed considering plant heights, *I. serracarajensis* presenting moderate growth with a gain of only 1 cm over the period evaluated, whereas *I. cangae* presents an approximately 3 cm growth during the period evaluated (Figure 3). *I*. *serracarajensis* are too short of reaching above the waterline. In the field, these plants are resistant to suddenly dry.

On the other hand, *I*. *cangae* only survives in one lagoon, always submerged. However, during *ex-situ* cultivation, *I*. *cangae* leaves were able to survive above the waterline. Here, we first report this remarkable trait. We experimented with reducing the water suddenly, but leaves shrink in a few minutes.

The leaf growth rate of *I. cangae* was 0.191 cm per day, while *I. serracarajensis* was 0.042 cm per day. These results are in confidence with biochemistry differences between the species, while *I. cangae* plants presented higher plasma membrane H^+^-ATPase activity as compared to *I. serracarajensis* (Figure 4). This enzyme plays a central role in the majority of plant processes, such as tissue expansion, nutrient acquisition, stress tolerance, and others (Falhof et al. 2016). The leaf elongation correlates to higher values of H^+^-ATPase activity (Figures 2 and 3).

**Figure 4.**
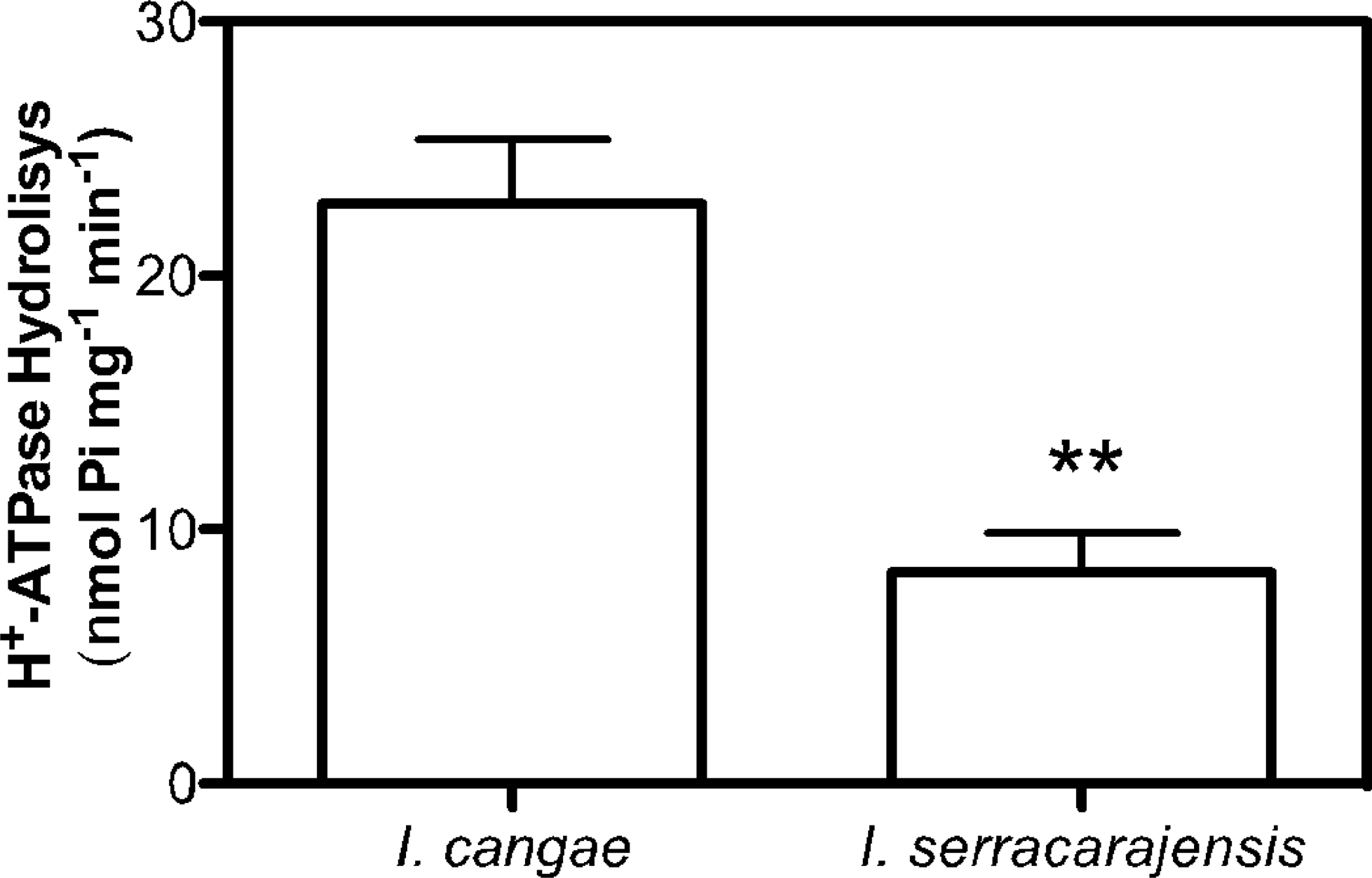
Plasma membrane H^+^-ATPase activity. Values are presented as means ± standard deviation (SD) of ten individual plants per specie; two asterisks indicate values that were determined by the t-test to be significantly different (P < 0.001).

Photosynthetic performance of the two *Isoetes* species evaluated thought the PSII efficiency was also distinct, *I. serracarajensis* PSII efficiency is near to 20% higher the *I. cangae* (Figures 5 and 6). *I. cangae* is only found submerged in permanent lagoons over “canga” soil, while *I. serracarajensis* is also found terrestrial on wet soil in addition to submerged (Pereira et al., 2016). Chlorophyll *a* fluorescence is considered a useful indicator of light reactions of photosynthesis (Rascher et al., 2000) and the photochemical efficiency of photosystem II can be assessed by PAM fluorometers (Schreiber et al., 1975). Data obtained have no precedence in the bibliography. To the best of our knowledge, the only work addressing PSII efficiency in *Isoetes* genus was published by Hawes and collaborators (Hawes et al., 2003). Herein we present the first report of PSII efficiency of these two new Amazon quillworts.

**Figure 5.**
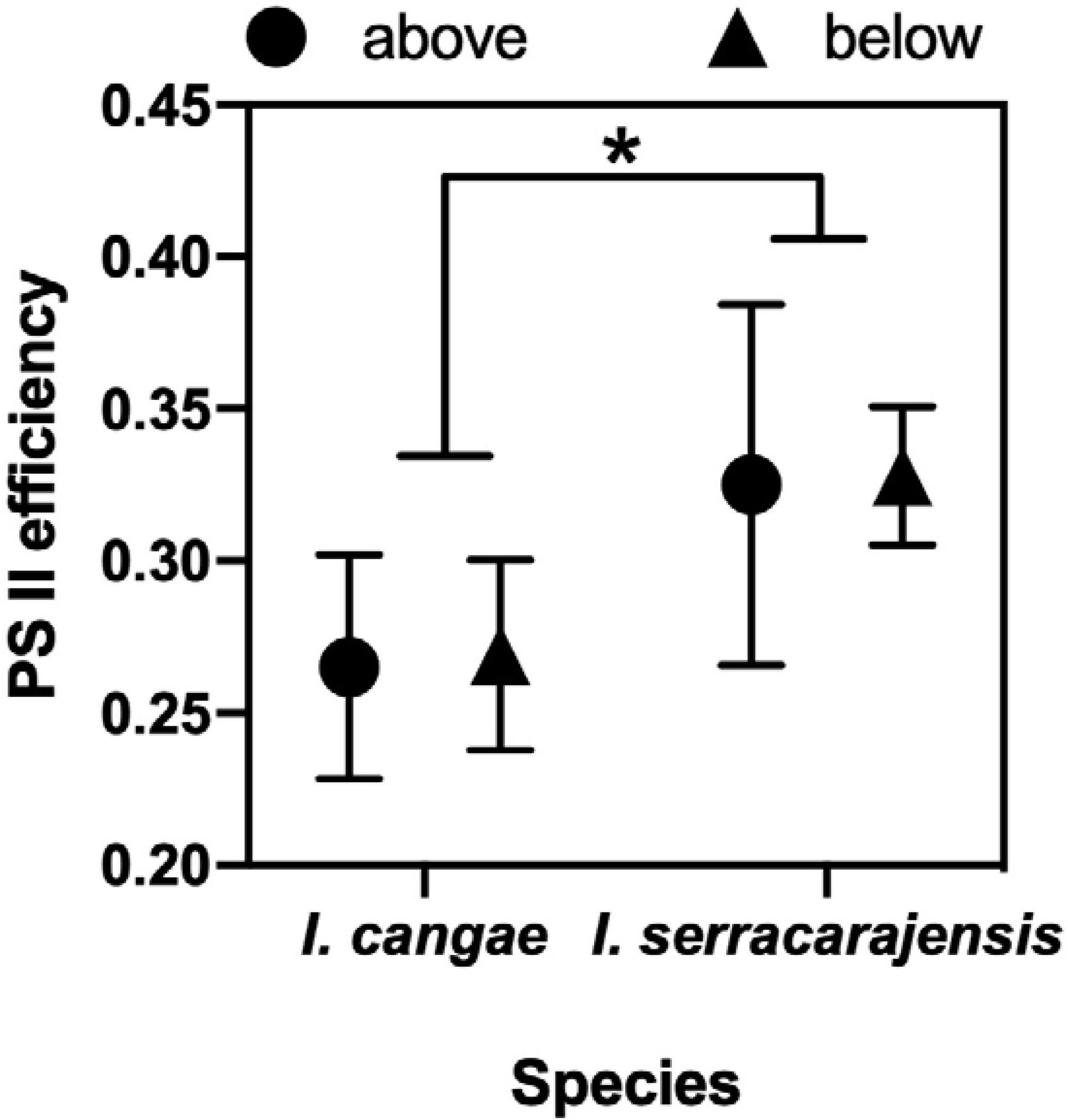
Photosynthetic performance of the two studied *Isoetes* species in regard to leaf exposition to air (black circle denotes leaves above water line, and black triangle denotes leaves below water line). The effective quantum yield of photosystem II (PS II) (Δ*F*/*Fm’*) was accessed on the two species at two leaf positions: below and above the water line. Values are presented as means ± standard deviation (SD) of ten individual plants per species. Asterisk denotes statistical difference between genotypes at P<0.05.

**Figure 6.**
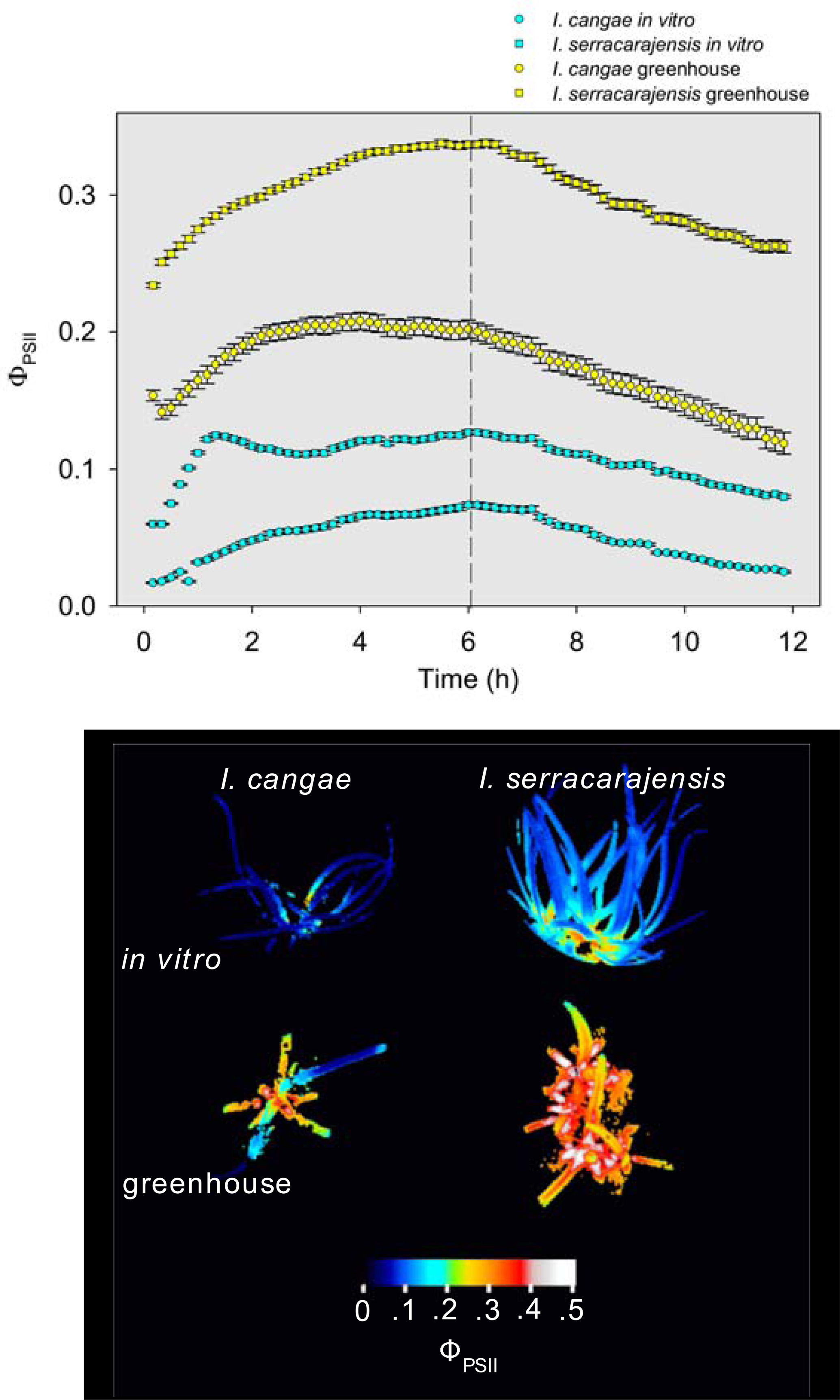
Diurnal course of Φ_PSII_ in two *I. cangae* and *I. serracarajensis* cultivated *in vitro* and in greenhouse obtained by serial imaging of Φ_PSII_ every 10 minutes within 12 hours. Plots are entire plant mean of Φ_PSII_ and error bars are confidence intervals at p<0.001 (Student’s t distribution). The image bellow show example plants measured at 6^th^ hour.

The leaf position to the waterline did not interfere in the PSII efficiency (Figure 5). The PSII efficiency of *I. serracarajensis* is near to 20% higher than the *I. cangae.* The values were 0.33 in *I. serracarajensis* and 0.27 in *I. cangae*. The results are hard to compare because our plants were cultivated *ex-situ* at 12 cm water depth, while *I. alpinus* plants were measured *in-situ* in 3-7 m depth. However, our results were in the range of 0.02 to 0.80 reported by (Hawes et al., 2003).

Different concentrations of iron (FeEDTA 0.20; 0.60 and 1.80 mg/L) than the observed in natural conditions (0.55 mg/L of iron in the habitat of *Isoetes cangae*) did not alter the growth parameter as measured by sprouting and leaf length (Figure 8). The sprouting rate was 0.089 per day on average in the different Fe concentrations. The leaf growth rate was 0.13 cm per day on average in the different iron concentrations.

**Figure 7.**
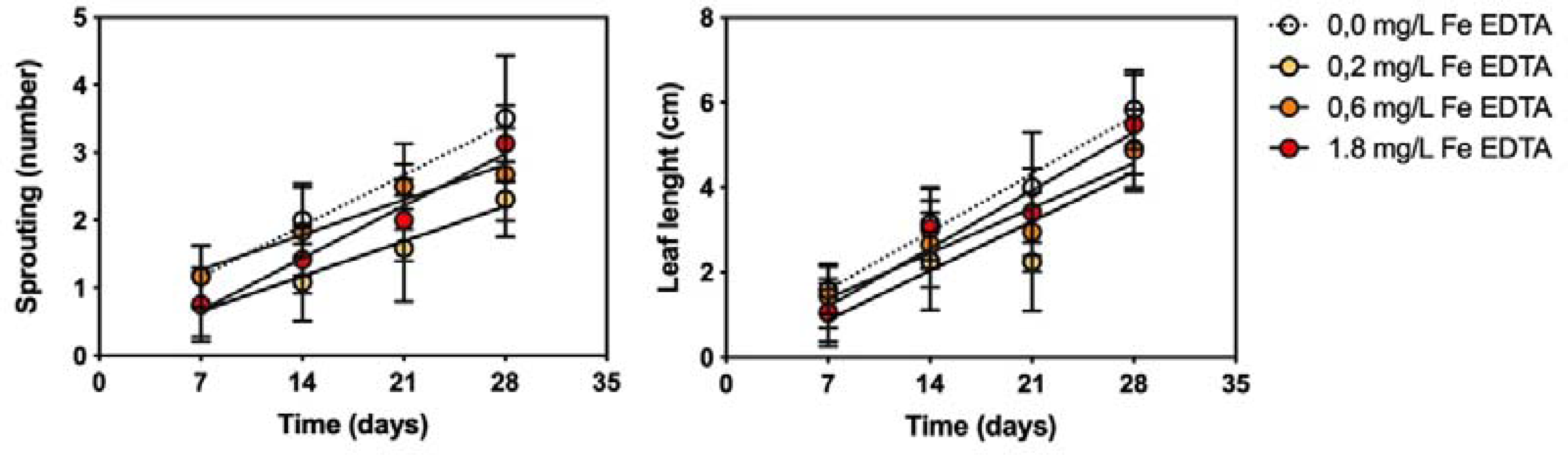
Sprouting rate and growth of *I. cangae* as a function of different concentrations of iron (FeEDTA 0.2; 0.6 e 1.8 mg/L). Dotted lines are linear regression of control plants (FeEDTA 0.0 mg/L).

**Figure 8.**
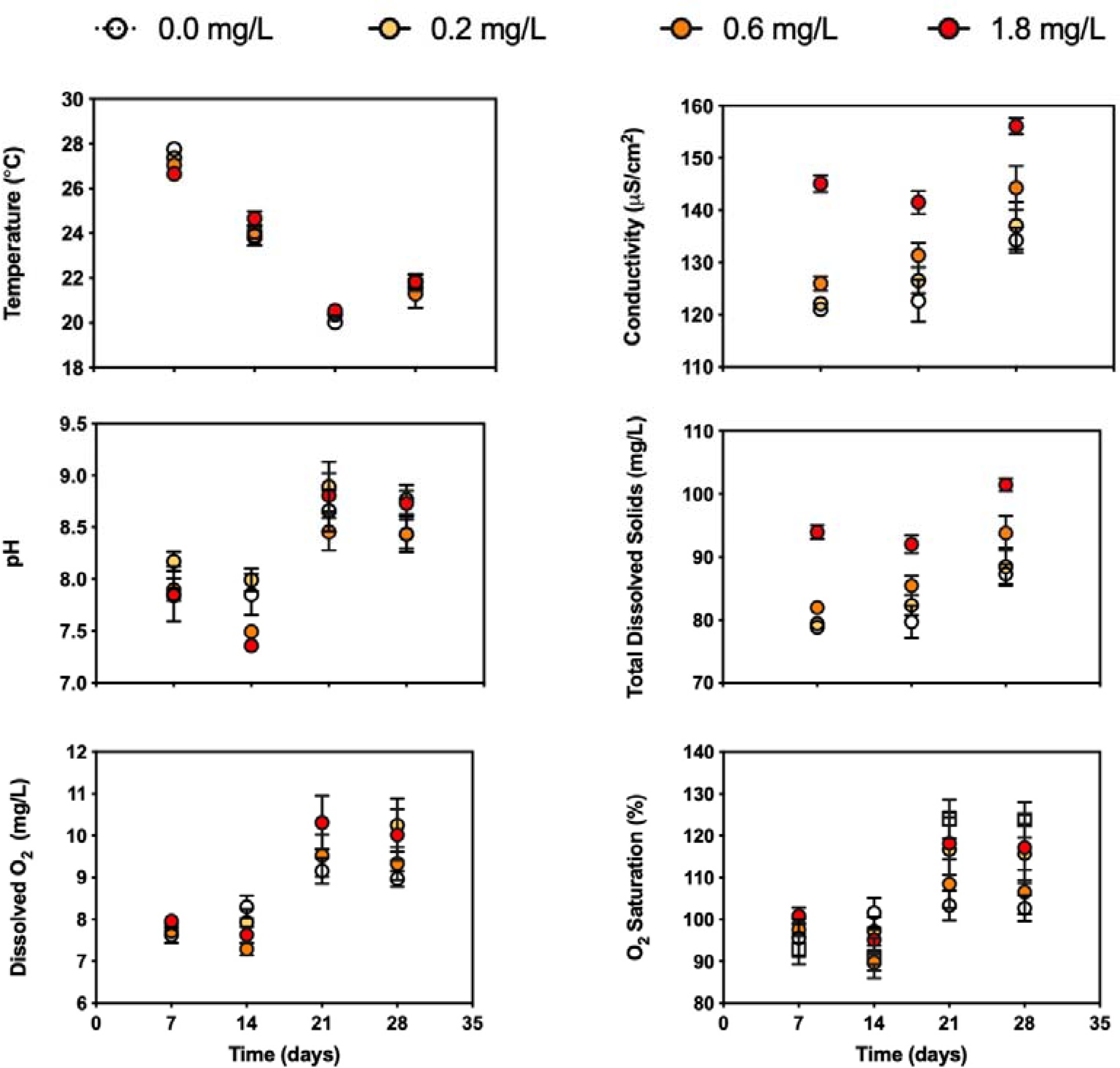
Physicochemical characteristics of the water in the beckers with different iron concentrations (FeEDTA 0.2; 0.6 e 1.8 mg/L). Temperature, pH, Conductivity, Salinity, Total Dissolved Solids, Dissolved O_2_ and O_2_ Saturation.

The sporophytes derived from *ex-situ* plants were able to growth in an artificial pool in Serra dos Carajás and near to 80% of plants survived planting in the lagoon after four months.

## 4. DISCUSSION

Generally, the water physicochemical characteristics in natural habitats and beakers of both species were very similar, but some parameters varied between the species. These parameters seem to be a critical started point to understand the ecology and physiology of *Isoetes*. A study reported the water chemistry and the distribution of the endangered aquatic quillwort *lsoetes sinensis* in China as a determinant for the distribution and occurrence of that species (Wen et al., 2003). However, the definition of a typical isoetid environment it is not quite simple (Boston and Adams, 1987; Smolders et al., 2002), because of the inherent geochemistry difference of the sites.

Values of total N, B, Al, P, Ca, Cu, Mn, Zn, and S in the lagoon of *Isoetes cangae* are very similar to the ponds of *Isoetes serracarajensis*. However, the ponds are more abundant in K (120-fold), Fe (2.4-fold), and Ca (2.0-fold), and more deficient in Na (40% less). Also, the pond water has a 6-fold higher electric conductivity and a slightly higher pH than the lagoon of *I. cangae*. Wen et al. (2003), studying aquatic quillwort *lsoetes sinensis* in China, reported higher values of total P, Ca, Mg, Mn, and Zn. Higher levels of water Ca and Mg was also observed in lagoons from North America with *Isoetes macrospora* (Boston and Adams, 1987). We have reported here a higher level of iron dissolved as expected due to the vast iron ore deposits of our study site (Gagen et al., 2019). Isoetids are found from sandy, low-nutrient, and low-organic sediments to high-organic, clay-rich sediment (Sand-Jensen and Søndergaard, 1978; Wilson and Keddy, 1985). Here the sediment of original sites of each species was characterized. and the values of P, Na, B, Co, Fe, Mn, and Zn were higher in the pond as compared to the lagoon (Table 3). Instead, organic matter, K, Ca, and Mg were higher in the lagoon. The pond dries seasonably, corroborating for the lower content of organic matter. Additionally, the lagoon has organic inputs from sources composed primarily of palms and macrophytes (Sahoo et al., 2016). The enormous Fe concentration is a reflex of the geochemistry of this enriched Fe_2_O_3_ sediment.

This work demonstrated that a simple cultivation system could be applied for *ex-situ* conservation for both *Isoetes cangae* and *Isoetes serracarajensis*. We do not find differences in plant growth on different seasons; however, we register variations on PAR average values, so *Isoetes* plants appeared to adapt very quickly to radiation changes. The pH of water of *I. cangae* plants was lower as compared to the *I. serracarajensis*′ at the begging of the experiments. At the end of the monitoring, the water of both species presented a similar pH. Very little is known about *Isoetes* habitat in the tropics, but they were found in water with a wide pH range from 5.1 to 8.51 in Europe, North America and Asia (Abeli et al., 2012; Boston and Adams, 1987; Chappuis et al., 2011; Wen et al., 2003). The O_2_ parameters of the water were initially higher in beckers of *I. serracarajensis* than those of *I. cangaès.* The majority of the literature does not present O_2_ data. The dissolved O_2_ in the water had varied from 6.07 to 7.47 mg/L in four study sites of *Isoetes sinensis* in China (Wen et al., 2003). The dissolved O_2_ in the water of the original site of *I cangae* is 7.75 mg/L on average (data not shown).

Since the organic matter did not appear to be constraint and it was previously used for *in vitro* cultivation of *Isoetes cangae* (Caldeira et al., 2019), we have used an organic substrate based on sphagnum peat and covered with sand that supplied plants effectively (Table 3). Generally, macronutrients were higher in the artificial substrate, while micronutrients were lower in the substrate.

Plants of *I. cangae* from the field presented tissue nutrient concentrations higher than plants of *ex-situ* cultivation system overall, except for Ca and B (table 2). The accumulation of metals such as Cu, Fe, and Mn in both species in the natural environment is notably higher than an artificial system. Considering soluble Fe concentration in the water, the most contrasting element concentration between habitats of isoetids plants described in bibliography and *I. cangae* e *I serracarajensis*, the soluble iron had not influenced the growth parameter as measured by sprouting and leaf length. However, iron concentrations in plant leaves are remarkably high. To the best of our knowledge, no other plant was ever described with such Fe accumulation capacity. *I. cangae* was able to accumulate 16,950 mg/kg total Fe and *I. serracarajensis* 33,050 mg/kg total Fe. *Isoetes anatolica* from seasonal ponds of Turkey can accumulate 431.00 mg/kg of Fe (Ozyigit et al., 2013). Here plant macronutrients were also evaluated, and *I. serracarajensis* presented more N, P, K, Ca, Mg, and S than *I. cangae*. It is rare data of plant nutrition in *Isoetes.* It was reported in *Isoetes lacustris* from the Lagoon Baciver in Spanish Pyrenees a range of 2.0 - 2.6% N and 0.14-0.26 % P (Gacia and Ballesteros, 1994). *I. lacustris* from Loch Brandy, Scotland, presented 1.8 % N and 0.02 % P (Richardson et al., 1984).

The leaves of *I. cangae* grown faster than *I. serracarajensis*. The leaf elongation might be associated with higher values of H^+^-ATPase activity (Figures 3 and 4). The regulation of H^+^-ATPase activity is closely related to tissue elongation through acid growth (Dünser and Kleine-Vehn, 2015). Most of the data is related to land plants, but similar mechanisms have been shown in aquatic plants (Koizumi et al., 2011). Plasma membrane H^+^-ATPase plays a central role in the majority of plant processes, such as tissue expansion, nutrient acquisition, stress tolerance, and others (Falhof et al., 2016). Ion transport studies in aquatic plants are rare (Babourina and Rengel, 2010; Baur et al., 1996), and our findings may contribute to understanding part of the mechanisms relying on aquatic plant health and development.

Monitoring enzymes activity in field conditions is unpractical. For field assessment of Isoetes’s health, the use of photochemical efficiency is a more practical ecoindicator. Chlorophyll a fluorescence is considered a useful indicator of light reactions of photosynthesis (Rascher et al., 2000) and the photochemical efficiency of photosystem II (PSII) can be assessed by PAM fluorometers(Schreiber et al., 1975). The photosynthetic performance of the two *Isoetes* species was evaluated through the PSII efficiency. To the best of our knowledge, the only work addressing PSII efficiency in *Isoetes* genus was published by Hawes and collaborators (Hawes et al., 2003). Herein we present the first report of PSII efficiency of these two new Amazon quillworts. The results are hard to compare because our plants were cultivated *ex situ* at 12 cm water depth, while *I. alpinus* plants were measured *in-situ* in 3-7 m depth. However, our results were in the range of the values reported by Hawes et al., (2003).

The data presented illustrate that both species can grow and reproduce in a artificial low-cost media composed by commercial organic substrate, sand, and tap water. Here, we first report the *ex-situ* cultivation of *Isoetes* in outdoor conditions. It is presented a very feasible protocol, where critical physiological aspects of plant growth were monitored. The technics employed here and the characteristics evaluated during *ex-situ* cultivation are relevant for understanding both the ecological and physiological traits of *Isoetes*. The results of this work have broad applicability, assisting other low-cost *ex situ* cultivation studies as an essential strategy for isoetids conservation.

